# Mosquito population dynamics are shaped by interactions among larval density, temperature, and relative humidity

**DOI:** 10.1101/2024.06.08.598043

**Authors:** Nicole Solano, Emily C. Herring, Carl W. Hintz, Philip M. Newberry, Annakate M. Schatz, Joseph W. Walker, Gregory R. Jacobs, Craig W. Osenberg, Courtney C. Murdock

## Abstract

Understanding how variation in key abiotic and biotic factors interact at ecologically relevant spatial scales is crucial for predicting population dynamics, distributions, and abundances. This is especially true for vectors that transmit human pathogens. However, studies investigating the effects of environmental variation on vectors have typically investigated environmental factors in isolation or in laboratory experiments that examine constant environmental conditions that often do not occur in the field. To address these limitations, we conducted a semi-field experiment in Athens, Georgia using the invasive Asian tiger mosquito (*Aedes albopictus*). We selected nine sites that varied in impervious surface and vegetation cover to explore effects of natural variation in microclimate (specifically, temperature and relative humidity) on mosquitoes. We manipulated conspecific larval density at each site and repeated the experiment in the summer and fall to further increase the variability in temperature and relative humidity. We then evaluated the original design features (land cover, larval density, and season), and their interactions, on the mean proportion of females emerging, juvenile development time, size upon emergence, and estimated per capita population growth (i.e., fitness). We found significant effects of larval density, land cover, and season on all response variables, including a non-intuitive decrease in development time with increasing larval density in the fall. We repeated these analyses using the hypothesized microclimate drivers of these effects: temperature and relative humidity. In general, the model using the microclimate variables outperformed the model using land use and season, highlighting the roles of temperature and relative humidity and their interactions with density on mosquito traits and dynamics. Our study demonstrates that ignoring the interaction between variation in biotic (e.g., intraspecific competition) and abiotic (e.g., temperature and relative humidity) variables could reduce the accuracy and precision of models used to predict mosquito population and pathogen transmission dynamics, especially those inferring dynamics at finer-spatial scales across which transmission and control occur.

**ABSTRACT (Spanish):** Es crucial comprender cómo los factores abióticos y bióticos interactúan a escalas ecológicamente relevantes para predecir la dinámica, distribución, y abundancia de poblaciones. Esto es especialmente cierto para vectores que transmiten patógenos humanos. Sin embargo, la mayoría de estudios han considerado factores ambientales de forma aislada o en condiciones de laboratorio poco realistas. Para superar estas limitaciones, realizamos un experimento de semi-campo en Athens, Georgia, con el mosquito invasor *Aedes albopictus*. Manipulamos la densidad larval en nueve sitios que variaban en cobertura vegetal y superficie impermeable, y repetimos el experimento en el verano y otoño. Luego, evaluamos los efectos de la cobertura del suelo, la densidad larval, y la estación en la proporción de hembras adultas emergidas, el tiempo de desarrollo, el tamaño al emerger, y el crecimiento poblacional per cápita. Encontramos efectos significativos de todas las variables, así como fuertes interacciones entre estación y densidad de mosquitos, incluyendo una disminución no intuitiva del tiempo de desarrollo en el otoño con mayor densidad larval. Repetimos los análisis usando factores microclimáticas hipotetizados como impulsores de estos efectos: temperatura y humedad. En general, el modelo que incorpora variables microclimáticas superó al modelo basado en el uso de suelo y estación, evidenciando la importancia de la temperatura y la humedad, así como sus interacciones con la densidad larval, en las características biológicas y la dinámica poblacional de los mosquitos. Nuestros resultados demuestran que la interacción entre variación biótica (competencia intraespecífica) y abiótica (temperatura y humedad), es fundamental para mejorar la precisión de los modelos que predicen la dinámica poblacional de mosquitos y la transmisión de patógenos a escalas locales de transmisión y control.

## INTRODUCTION

Three processes generally determine the distribution and abundances of organisms: environmental filtering (abiotic variables), species interactions (biotic variables), and constraints on organismal dispersal (Leibold and Chase, 2018; Menge and Sutherland, 1976; Vellend, 2016). The effects of abiotic and biotic variables on species distribution and abundances have become some of the most important issues in ecology today, especially in the context of rapid biodiversity loss (McKinney and Lockwood, 1999) climate change (Parmesan, 2006) and human-mediated dispersal (Bullock and Pufal, 2020). Multiple environmental factors (both abiotic and biotic) affect the demography, vital rates (e.g., development, growth, and survival), and population dynamics of ectothermic organisms. However, predicting the effects of abiotic and biotic variables on organismal distribution and abundance patterns is difficult because their effects on processes that determine organismal fitness and population dynamics are scale-dependent (Levin, 1992; McGill, 2010; Wiens, 1989).

It has been long hypothesized that environmental filtering is strongest at larger, more regional scales because climate variables (e.g., temperature, relative humidity, precipitation) exhibit little variation at fine-spatial scales (Houghton et al., 2002; Levin, 1992; McGill, 2010; Wiens, 1989). With the emphasis on climate warming in the ecological literature, there has been a strong focus on understanding the effects of temperature on the distribution and abundance of ectothermic organisms, as well as their traits ( Helmuth et al., 2010; Mordecai et al., 2019; Parmesan, 2006). These effects have been leveraged in species distribution models and mechanistic models to predict the current and future geographic ranges of ectothermic organisms (Amarasekare, 2024; Huey et al., 2001; Simon and Amarasekare, 2024). In contrast to the large spatial scale for abiotic variables typically assumed in species distribution models, many abiotic variables are likely to exhibit variation at small-scales (e.g., contrasting temperatures between sunny and shaded microhabitats). For small organisms (e.g., annual plants, reptiles, insects), this local variation in abiotic conditions might produce variation in demographic traits that could not be anticipated from average conditions across larger spatial scales. However, the effects of this small-scale variation in abiotic conditions is often ignored.

Biotic variables, like intra-and inter-species competition, availability of resources, and presence and density of biological enemies (predators and parasites), also have important effects on organismal fitness and population dynamics (Beale et al., 2014; Golding et al., 2015; Leach et al., 2016). These biotic factors affect individuals due to interactions within their local neighborhood; thus, the effects of these factors on species distribution and abundance patterns are hypothesized to arise at fine-spatial scales (Levin, 1992; McGill, 2010) but are often not incorporated into species distribution or mechanistic models (Cohen et al., 2016).

Finally, abiotic and biotic factors can interact to affect species distributions, which could be especially relevant for organisms with complex life cycles, such as amphibians or insects, that use very different habitats across their life stages (Govindarajulu and Anholt, 2006; Indermaur et al., 2010). When spatial variation in these biotic and abiotic variables is ignored, predictions of a given species’ population dynamics averaged over space or time can lead to considerable bias if the effects of these factors are nonlinear (Bozinovic et al., 2011; Martin and Huey, 2008; Ruel and Ayres, 1999). As a result, mathematical models might overlook the influence of particularly advantageous microsites (e.g., hot spots) or deleterious microsites (e.g., ecological traps) on a given species’ population dynamics.

One of the areas in which these issues are readily apparent is in the field of mosquito ecology. Climate and land use change are rapidly altering the landscapes across which mosquitoes occur, altering the ecological relationships mosquitoes share with their environments and their vertebrate hosts. This has led to biological invasions of mosquito species and their parasites to new areas of the world (Roiz et al., 2024; Taylor et al., 2024) with consequences for the emergence of new diseases (Chala and Hamde, 2021; Lounibos, 2002; Yan et al., 2024). Understanding and predicting how future changes in climate and land use will alter mosquito dynamics and the spread of vector-borne pathogens is particularly relevant. Thus, there has been considerable investigation into the effects of temperature on mosquito distributions, population dynamics, and parasite transmission (reviewed in Kraemer et al., 2019; Ryan et al., 2023; Sinka et al., 2020). Recent work also suggests that relative humidity can have important effects on larval traits and mosquito population dynamics, possibly through effects of relative humidity on the surface tension of habitats (see Brown et al., 2023; Huxley et al. 2025). In addition to abiotic variables, mosquitoes experience intra-and inter-specific competition during their larval stage, which can affect mosquito survival, reproduction, and generation time by slowing development, reducing adult size, and lowering teneral reserves (e.g., lipids and proteins) of adult mosquitoes (Briegel, 2003) with significant implications for mosquito fitness and vectorial capacity (Alto et al., 2005; Moller-Jacobs et al., 2014; Shapiro et al., 2016).

Currently, there has been little investigation of the potential interactive effects of abiotic and biotic factors on mosquito population dynamics. Most studies that have considered the interaction between abiotic (i.e., temperature, relative humidity) and biotic factors (i.e., intraspecific competition) on mosquito life history traits and population dynamics have been conducted in the lab under constant environmental conditions that may not reflect field environments (Evans et al., 2021; Tesla et al., 2018; Shapiro et al., 2017). In the field, abiotic variables like temperature vary diurnally, seasonally, and across space (e.g., due to variation in land use and land cover) (Evans et al., 2019; Murdock et al., 2017; Wimberly et al., 2020).

To improve our understanding of how field-relevant variation in key abiotic and biotic factors interact at spatial scales relevant for mosquito fitness and population dynamics, we conducted a semi-field experiment using the Asian tiger mosquito (*Aedes albopictus*) in Athens, Georgia during summer and fall 2017. *Ae. albopictus* is a highly invasive mosquito in the United States and other areas of the world, as well as a competent vector of arboviruses including dengue and chikungunya (Benedict et al., 2007; Bonizzoni et al., 2013). We selected nine sites that varied in impervious surface and vegetation cover, and conducted an experiment across the sites at two different times to leverage natural spatial and temporal variation in temperature and relative humidity to explore their effects on mosquito traits and population dynamics (Brown et al., 2023; Murdock et al., 2017; Wimberly et al., 2020). During each experiment, we manipulated conspecific larval density, a biotic variable known to have large effects on metrics of mosquito fitness and population growth rates (Juliano, 2009; Teng and Apperson, 2000). We simultaneously monitored spatial and temporal variation in temperature and relative humidity. Although we designed our study based upon density, land use and season, we were specifically interested in the interactive effects of conspecific density, temperature and relative humidity, each of which is known to affect mosquito demography in isolation (e.g., Alto and Juliano, 2001; Brown et al., 2023; Mordecai et al., 2019), while controlling for other factors that might vary with season and land use (e.g., leaf litter or detrital inputs). We hypothesized that temperature and relative humidity would serve as “master variables” and would provide a more general and better predictor of mosquito dynamics than land use and season. Understanding the drivers of spatial and temporal heterogeneity on the dynamics of invasive species like *Ae. albopictus* is crucial for predicting and managing their future invasive potential, as well as for other species with similar ecology (e.g., *Aedes aegypti)* that play an important role in human arbovirus transmission.

## METHODS

### Experimental design

We conducted the study between June 2017 and October 2017 in Athens-Clarke County, Georgia, USA. We utilized nine previously established field sites that ranged in impervious surface, vegetation cover, and local microclimate (Appendix S1: Figure S1) (Evans et al., 2019). Within each of the nine sites, we placed three trays in full shade separated by approximately 6-9 m. Each tray held four wide-mouth mason jars each containing 300 mL of leaf infusion (Evans et al., 2018; Murdock et al., 2017). Three jars in each tray received a specific larval density treatment (30, 60, or 120 *Ae. albopictus* first instar larvae from a laboratory colony established from field collected *Ae. albopictus* in 2015, Athens, GA; Murdock et al., 2017). We covered the top of the jars with mesh netting and staked a wire cage with a vinyl roof over the trays (see Murdock et al., 2017) to prevent changes in habitat volume from rainfall, the addition of nutrients due to falling vegetation, and the loss of jars from animal disturbance. To understand how seasonal variation in abiotic factors may alter relationships between conspecific larval density and traits governing mosquito population dynamics, we conducted this experiment in the summer (June 11 - July 25) and fall (September 8 - October 18) 2017. Jars were visited daily and at the same time of day. Any adults that had emerged were collected and placed on ice for transport back to the lab. Mosquitoes were then stored at-20℃ until we determined their sex (Darsie and Ward, 2005) and wing length.

### Microclimate variables

To measure variation in water temperature, we deployed one data logger (Monarch Instruments, Amherst, NH, USA: Radio Frequency Identification (RFID) Temperature Track-It Logger) in one jar in each tray that had been filled with leaf infusion but contained no larvae. We also uniformly distributed six data loggers of the same brand throughout each site at approximately 0.5-1.0 m above the ground and in full shade to measure relative humidity. All loggers recorded microclimate variables every ten minutes with data downloaded monthly.

### Mosquito life history traits

In this study, we only report female data since females determine mosquito population growth rates and pathogen transmission (Powell, 2025). To calculate the proportion of mosquito larvae and pupae that survived to emerge as adults (i.e., emergence), we divided the total number of females that emerged by half the total number of 1^st^ instar larvae placed into the habitat (e.g., 15, 30, or 60 assuming half of the 1^st^ instar larvae were female: Lounibos and Escher, 2008). Juvenile development time was defined as the number of days between the start of the experiment and the day an adult female emerged. One wing was removed from each individual, mounted onto a glass slide using clear nail polish, and measured with a dissecting microscope and micrometer to index body size (Packer and Corbet, 1989). To estimate the per capita growth rates of mosquito populations in response to our experimental treatments we used equation 7 from Livdahl & Sugihara (1984):

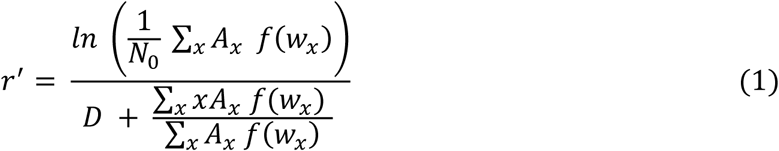

where *N*_0_ is the initial number of females in a cohort (i.e., half of the density treatment), *A_x_* is the number of females that emerged on day *x*, and *D* is the time to reproduction following emergence (which we assume to be 14 days: Lounibos et al., 2002). The fecundity *f(w_x)_* of females that emerged on day *x*, with an average wing length of *w_x_*, was estimated from the linear relationship between wing length and number of eggs laid from Lounibos et al. (2002):

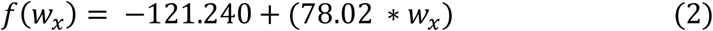

In 12 cases, no females emerged from a jar (these all occurred in the fall at high density). Since removing those jars would elevate and bias *r*^′^ for that treatment, we imputed their per capita growth rate by setting *r*^′^ =-0.05 day^-1^, which was just below the most negative *r*^′^ observed in the other jars.

### Data Analyses

We took three complementary approaches to analyze the results. First, we examined variation in water temperature and relative humidity across land use and season. To assess how daily microclimate conditions varied, we used linear mixed effects models to investigate how the daily mean, minimum, maximum, and range in air temperature (T_mean,_T_min,_T_max,_T_range)_and relative humidity (R_mean,_R_min,_R_max,_R_range)_varied with design variables. All models included fixed effects of season, land-use type, and their interaction, and random effects of site, logger (nested within site), and day.

Second, we investigated how juvenile survival and development rate, adult size, and estimated per capita population growth (i.e., fitness) of *Ae. albopictus* varied with land use, season, and larval density. Although our putative interest was microclimate (and not land use and season per se), our sampling design was based on these categorical factors, so we started with those analyses before moving on to exploration of the effects of temperature and relative humidity. We also used GLMMs to determine how emergence (binomial distribution; *logit* link function), larval development rate (lognormal distribution; *log* link function), wing length (lognormal distribution; *log* link function), and per capita growth rate (Gaussian distribution; *identity* link function) varied across seasons, land-use types, and density treatments. Those generalized linear mixed effects models all had season, land-use type, density treatment, and their interactions as fixed effects, with site and tray (within site) as nested random effects. Analyses of development rate and wing length also included jars (within tray) as an additional nested random effect.

Third, we constructed models that replaced land use and season (the design variables) with temperature and relative humidity (the putative proximate drivers of variation mosquito dynamics). To begin our analyses of the effects of microclimate on mosquito traits, we first explored the correlations among the microclimate variables to avoid multicollinearity. Based on correlation analyses, we included only T_min,_T_max,_and R_min i_n our models since they were of particular biological importance to us and they were not strongly correlated with each other (Appendix S1: Figure S2). We analyzed effects of the microclimate variables using GLMMs and the same distributions and link functions for each response variable as described above, although in this case the predictors were continuous rather than categorical.

Finally, to compare the use of the microclimate variables vs. season and land use for each response variable, we ran three models – *null*, *base*, and *microclimate* – and used the corrected Akaike Information Criterion (AIC_c)_ to evaluate their relative performance. The *null model* only included density treatment as a fixed effect (e.g., it lacked land use and season). The *base model* incorporated the original design factors (season, land-use type, density, and their interactions) as fixed effects. The *microclimate model* replaced the effects of land use and season with the microclimate variables, and therefore included the fixed effects of density, along with the main effects of T_min,_T_max,_R_min a_nd their interactions with density. For each response variable, all three models included the effect of density and an identical random effect structure, but they varied in how they incorporated environmental variables. Microclimate variables experienced by larvae in a jar were characterized by the mean environment experienced by those larvae as determined by their distribution of emergence dates (i.e., for each larva we calculated the mean experienced microclimate per larvae from day 0 to its day of emergence and then averaged these means across all females that emerged from the jar). In 12 cases in the fall at high density, no females emerged from a jar, so the microclimate variables for those jars were imputed based on emergence patterns from other jars at that site, season and density. Two sites did not have any females emerge (one rural site and one suburban site); therefore, we used the emergence patterns from other jars at the two other sites of the same land-use type, season, and density.

All analyses were conducted in R 4.6.0 (R Core Team, 2026). Code to run analyses and create figures can be found on Figshare (Solano, 2026). Models were fit using the *glmmTMB* package (Brooks et al., 2017), and compared by AIC_c u_sing the *bblme* package (Bolker and R Development Core Team, 2023). Type III analysis of variance (ANOVA) was conducted for all models using the *car* package (Fox et al., 2024). The *emmeans* package was used to obtain means and confidence intervals after adjusting for random effects (Lenth et al., 2025). We assessed model fit using the scaled residuals from the *DHARMa* package (Hartig et al., 2024).

## RESULTS

### Microclimate

Five of the 54 aquatic temperature loggers failed (three in the summer and two in the fall), and 36 of the 108 terrestrial loggers failed or were stolen (16 in the summer and 20 in the fall). Despite these losses, failure rates were roughly equivalent across sites and microclimate variation across season and land use types was well represented by the remaining loggers. Water temperature (i.e., mean daily T_min,_ T_mean,_and T_max)_ and relative humidity (mean daily R_min,_ R_mean,_and R_max)_metrics all varied significantly with season, land use, and their interaction (Appendix S1: Figure S3, Tables S1 and S2), demonstrating that land use and season captured significant spatiotemporal variation in mosquito-relevant microclimates.

### Emergence

Of the 5670 first instar *Ae. albopictus* placed into the field during each season, a total of 3156 mosquitoes (56% of the starting population) successfully emerged as adults during the summer while only 1233 mosquitoes (38%) emerged in the fall (emergence is equivalent to survival through the larval and pupal stages). Assuming that the initial frequency of females was 50%, there were significant main effects of season and conspecific density on the proportion of female first instar larvae that survived to emergence, with greater survival in the low-density treatment relative to medium-and high-density treatments (Figure 1a, Appendix S1: Table S3). There was also an interaction between season and density (Figure 1a) as well as a marginally non-significant interaction between land-use type and density (Appendix S1: Figure S4, Table S3). For example, there was little difference in the survival of female juveniles in the low-density treatments in the summer and fall; however, the negative effects of increasing conspecific density on juvenile survival was much greater in the fall (Figure 1a). Further, female juveniles at low conspecific densities survived best in suburban sites, while those at intermediate densities survived best in urban sites and those at high densities survived best in rural sites (Appendix S1: Figure S4).

**Figure 1.**
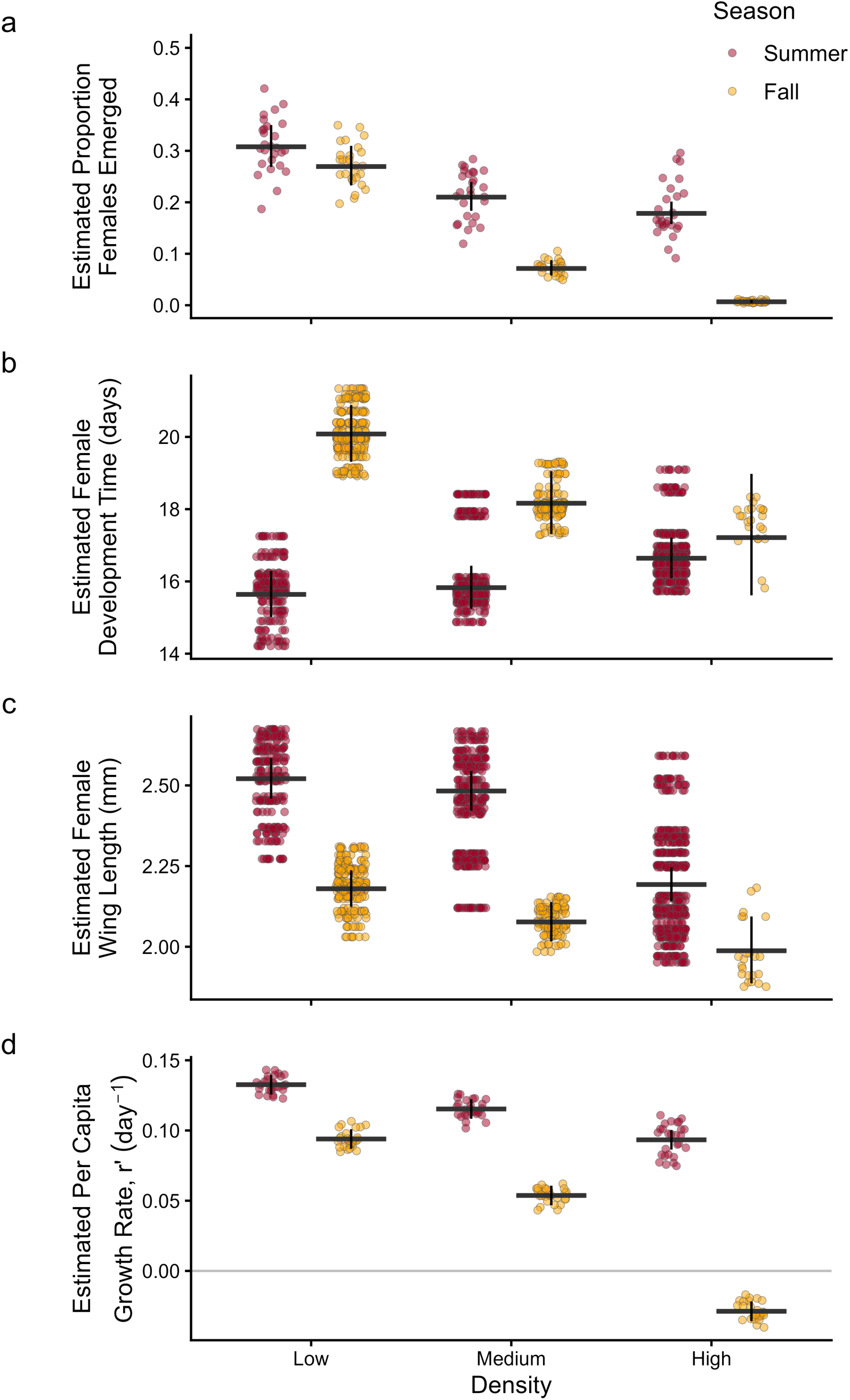
Results of mixed-effects model for *Ae. albopictus:* (**a**) the proportion of female larvae that emerged (i.e., survival probability), (**b**) female development time (days), (**c**) female wing length (mm), and (**d**) estimated per capita growth rate, *r*′ (day^-1^), for each density treatment (low = 30 larvae, medium = 60 larvae, high = 120 larvae) across two seasons (summer = red, fall = orange). Horizonal and vertical lines give the mean +/- 95% CI, and the dots indicate raw values [based on females in (b), (c), and jars for (a), (d)] adjusted for random effects.

Some of these effects likely arose through the influence of the microclimate variation across land use and season. For example, the microclimate model substantially outperformed the null model (ΔAIC_c =_393; Table 1), with the probability of female emergence (juvenile survival) decreasing with increasing density and increasing with increasing temperature (T_min)_ and relative humidity (R_min)_. In addition to these main effects, we observed interactions of both microclimate variables with density (Figure 2a): as density increased, the positive effect of temperature (i.e., T_min)_and relative humidity (R_min)_increased in magnitude (Figure 2a). However, the base model outperformed the microclimate model (ΔAIC_c =_6.3; Table 1), suggesting that the temperature and humidity variables did not capture as much of the spatiotemporal drivers of juvenile survival as did the original design variables (land use and season).

**Figure 2.**
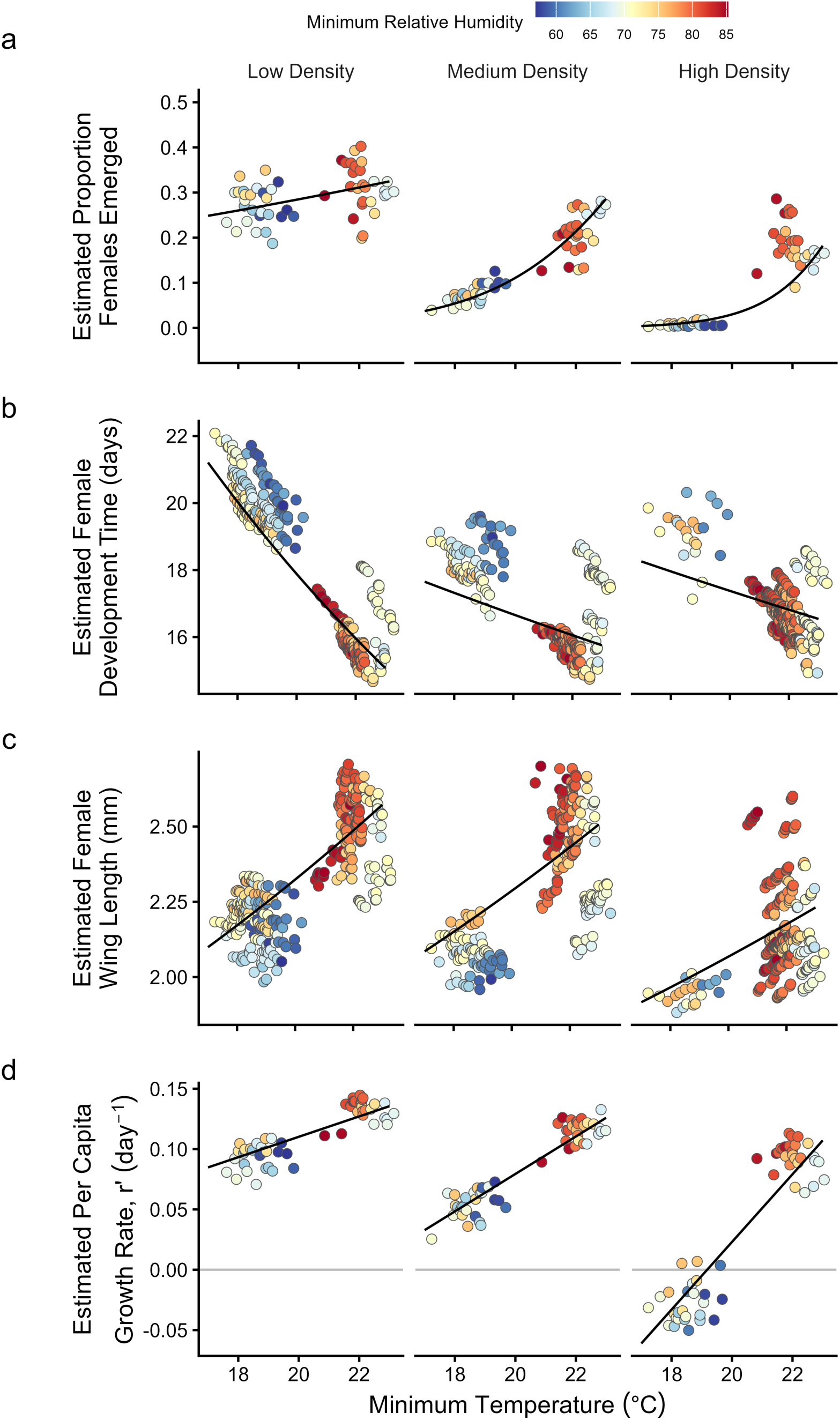
Results of mixed-effects model for *Ae. albopictus:* (**a**) the proportion of female larvae that emerged (i.e., survival probability), (**b**) female development time (days), (**c**) female wing length (mm), and (**d**) estimated per capita growth rate, *r*′ (day^-1^), for each of the three density treatments (low = 30 larvae, medium = 60 larvae, high = 120 larvae), as influenced by minimum temperature (T_min,_°C), and minimum relative humidity (R_min,_%). Dots indicate raw values [based on females in (b), (c), and jars for (a), (d)] adjusted for random effects. Black lines give the predicted effects of T_min f_or each density treatment.

**Table 1.** Results from Generalized Linear Mixed Effects Models testing the effects of larval density treatment (30 = low, 60 = medium, 120 = high), land-use type (rural, suburban, urban), and/or microclimate (T_min,_T_max, Rmin)_on mosquito traits across summer & fall seasons. The mosquito traits were analyzed using three model structures: (1) the Null model – main effect of larval density (but no other fixed effect); (2) the Base model – main effects of season, land use, and larval density and their interactions; and (3) the Microclimate model – main effects of larval density, water temperature (T_min,_T_max)_and relative humidity (R_min)_and the interactions between density and T_min,_ T_max,_and R_min._ Experimental site and tray (nested in site) were included as random factors in all models, and jar (nested in tray) was included for development time and wing length models. ΔAIC_c v_alues, k (number of parameters in the model), and statistically significant fixed effects from Type III analysis of variance tables are shown. Significance, based on the Wald test, is denoted as *p < 0.05* (*), *p < 0.01* (**), and *p < 0.001* (***). The best model(s) based on ΔAIC_c ≤_ 2 is indicated in bold. Full results for each model are provided in Appendix S1, Table S3.

| Response Variable | Model | $\Delta AIC_c$ | k | Significant Predictors |
| --- | --- | --- | --- | --- |
| Emergence | Null | 399.3 | 5 | Density*** |
|  | **Base** | **0** | **20** | **Season\*\*, Density\*\*\*, Season x Density\*\*\*** |
| | Microclimate | 6.3 | 14 | Density***, $T_{\min}$***, $R_{\min}$***, Density x $T_{\min}$***, Density x $T_{\max}$*, Density x $R_{\min}$*** |
| Development Time | Null | 92 | 7 | Density* |
|  | Base | 30.5 | 22 | Season***, Density*, Season x Density*** |
|  | **Microclimate** | **0** | **16** | **Density\*, $T_{\min}$\*\*\*, Density x $T_{\min}$\*** |
| Wing Length | Null | 80.4 | 7 | Density*** |
|  | Base | 18.7 | 22 | Season***, Density***, Season x Density* |
|  | **Microclimate** | **0** | **16** | **Density\*\*\*, $T_{\min}$\*\*\*, Density x $R_{\min}$\*** |
| Per Capita Growth Rate (r') | Null | 215.4 | 6 | Density*** |
|  | **Base** | **0** | **21** | **Season\*\*\*, Density\*\*\*, Season x Density\*\*\*** |
|  | **Microclimate** | **1.1** | **15** | **Density\*\*\*, $T_{\min}$\*\*\*, $R_{\min}$\*\*\*, Density x $T_{\min}$\*\*\*, Density x $R_{\min}$\*\*\*** |

### Female Development Time

Both season and conspecific larval density significantly affected the average amount of time required for adult female emergence (Table 1, Appendix S1: Table S3), with females on average developing more quickly in the summer (day 9 mean emergence date) relative to the fall (day 13 mean emergence date) and in the low-density treatment (Figure 1b, Appendix S1: Figure S4b). However, a significant interaction between season and larval density demonstrates that the main effect of density is being driven by the relationship between mean female development time (time to emergence) and larval density in the summer (Appendix S1: Figure S4, Table S3). Females reared in the low-density treatment were the fastest to develop in the summer but were the slowest to develop in the fall (Figure 1b).

These effects on development time were better predicted using the microclimate variables (ΔAIC_c >_ 30 for the microclimate model). Development time decreased with increasing daily T_min,_ and this effect was strongest for mosquitoes developing in low density environments (Figure 2b). In contrast to the other life history parameters, there was little evidence to suggest a role of relative humidity on development time (Table 1).

### Wing Length

We observed significant main effects of season and density on female wing length (Appendix S1: Figure S4, Table S3). Females reared in the summer had longer wings compared to the fall, and females developing in the high conspecific larval treatment had significantly shorter wings than mosquitoes from the low-density treatment (Figure 1c, Appendix S1: Figure S4). The observed variation in wing length was better predicted by the microclimate model compared to the null or based models (Table 1: ΔAIC_c >_ 18). Increased T_min w_as associated with increased wing length (Figure 2c, Table 1). Additionally, we observed a significant interaction between density and daily R_min,_with increases in relative humidity resulting in increased wing size at low and medium densities but not at high density (Figure 2c, Table 1).

### Estimated Per Capita Population Growth Rate

Integrating these responses revealed strong effects of season, conspecific larval density, and the interaction between these two factors on the estimated per capita population growth rate, *r*′ (Equation 1, Figure 1d, Appendix S1: Table S3). In general, the estimated per capita growth rate was greater in the summer relative to the fall and was greater at low densities than at high densities (Figure 1d). A significant interaction between season and density was observed due to greater declines in the estimated per capita growth rate with increasing conspecific density in the fall relative to the summer (Figure 1d, Appendix S1: Tables S3). Additionally, we observed negative per capita growth rates at the highest conspecific density in the fall, while all other combinations of factors led to positive projected per capita growth (Figure 1d, Appendix S1: Figure S4). These effects may have arisen through the effects of T_min a_nd R_min,_both of which were positively associated with increased per capita growth (Figure 2d, Table 1), although both microclimate variables also exhibited significant interactions with density. For example, the effect of increasing T_min w_as greater on larvae living at high densities and more muted for those living at lower densities (Figure 2d).

### Effects of microclimate on mosquito performance

In presenting our results, we initially emphasized the effects of our base model because our study was designed with respect to land use and season. However, for two of the four response variables (development time and wing length), the model based on microclimate performed better than the base model by a considerable degree (e.g., ΔAIC_c >_ 18: Table 1), and the microclimate and base models performed similarly for per capita population growth rate (ΔAIC_c=_1.1). In contrast, the base model performed better than the microclimate model for emergence (ΔAIC_c=_6.3). The null model, which excluded abiotic variables, always performed the poorest (Table 1; ΔAIC_c >_80). In general, the microclimate models showed that mosquito performance increased with increasing warming temperature (T_min o_r T_max)_and increasing minimum relative humidity (R_min)_, although the magnitude of these effects often depended on density (Table 1, Figure 2). Thus, the microclimate effects captured the observed variation in mosquito traits as effectively as season and land use and further highlighted the interactions of biotic (density) and with abiotic (microclimate) variables.

## DISCUSSION

To date, the majority of studies investigating the effects of various abiotic and biotic factors on mosquito fitness, population dynamics, vector competence, and transmission potential have explored the effect of varying a single environmental variable such as temperature (e.g., Agyekum et al., 2022; Liu-Helmersson et al., 2014; Monteiro et al., 2007), conspecific density (e.g., Hawley, 1985; Reiskind and Lounibos, 2009; Sauers et al., 2022), or food resources (e.g., Levi et al., 2014; Telang and Wells, 2004). Further, these studies are often conducted in laboratory environments under relatively constant conditions. However, mosquito populations in the field will be subject to the effects of many different environmental variables that fluctuate diurnally and seasonally and likely interact. Previous work in the *Ae. albopictus* system suggests that subtle variation in microclimate in the field can translate into significant variation in mosquito life history traits, population dynamics, and arbovirus transmission potential (Evans et al., 2019; Murdock et al., 2017; Wimberly et al., 2020). Our experimental study builds on these results by showing that season and land use (via their effects on temperature and relative humidity) can interact with variation in key biotic variables (conspecific density) to affect metrics of mosquito fitness. The results from this study have broad implications for our understanding and ability to predict the effects of current and future environmental variation on mosquito fitness and population dynamics, and the environmental suitability for mosquito-borne disease transmission.

### Land use affects microclimate with subtle effects on metrics of mosquito fitness and population dynamics

Like previous studies conducted in the same region (Athens, Georgia) (Evans et al., 2018; Murdock et al., 2017; Wimberly et al., 2020), urban sites had higher minimum temperatures than rural sites and were drier than non-urban land use types. This is consistent with the Urban Heat Island (UHI) effect, where nighttime temperatures (analogous to T_min)_are 1-2.5°C higher (Hibbard et al., 2017) than surrounding areas due to the higher percentage of impervious surfaces in urban areas that absorb, store, and re-emit heat (e.g., asphalt and concrete). However, in non-urban areas, hydrologically permeable land covers re-emit less heat (e.g., forests, shrublands, wetlands), have higher evapotranspiration rates, and often provide ample shading resulting in lower temperatures and higher relative humidity than urban areas (Bowler et al., 2010; Brown et al., 2023; Huang et al., 1987). Despite the variation in microclimate observed across land use, this variation was small (Appendix S1: Figure S2) and for the most part did not significantly affect components of mosquito fitness or estimated per capita growth rates within a given season (Appendix S1: Figure S4). This is not necessarily surprising as Athens-Clarke County is a small city in Georgia, with narrower gradients of impervious surface cover, vegetation cover, and ambient temperatures than seen across geographically larger urban areas in the U.S. and worldwide (Oke, 1973; Peng et al., 2011; Zhou et al., 2017). We did observe effects of land use on the relationship between conspecific density and juvenile mosquito survival (i.e., emergence), suggesting the effects of intra-specific competition could potentially be modified across an urban gradient. However, while significant, the effects of the interaction between land use and density on metrics of mosquito fitness were relatively small (Appendix S1: Figure S4), likely because variation in microclimate (e.g., T_min a_nd R_min)_ was not large across Athens within a given season (Appendix S1: Figure S3).

### Seasonal variation in microclimate affects metrics of mosquito fitness and population dynamics

Seasonal variation in microclimate was much larger than what we observed across land use, with temperature and relative humidity being higher during the summer field trial relative to the fall. As a result, we observed strong seasonal effects on juvenile survival, the juvenile development time, adult body size, and estimated per capita growth rates. Mosquitoes are ectothermic, and their trait performance has been shown to have non-linear relationships with temperature variation across a diversity of mosquito species (reviewed by: Mordecai et al., 2019). For example, laboratory work conducted under a range of constant temperatures predicts *Ae. albopictus* juvenile survival and mosquito development rates are maximized at ∼25°C and ∼35°C, respectively (Mordecai et al., 2017). Thus, it is unsurprising that we generally saw faster juvenile development times and increased adult female emergence in the warmer summer field trial (with average temperatures ranging from 25-25.7°C) than the cooler fall (with temperatures ranging from 21.1-21.6°C) (Figures 1, 2, Appendix S1: Figure S3).

### Seasonal variation in microclimate shapes the relationship between intra-specific competition and metrics of mosquito fitness and population dynamics

In general, our results support previous research that shows decreasing temperature or increasing conspecific larval densities result in decreased juvenile survival, smaller adults upon emergence, and decreased per capita growth rates (Lord, 1998; Sauers et al., 2022; Walker et al., 1991). However, our results also highlight a critical and novel role of seasonality and land use in modifying the effects of conspecific larval density on mosquito fitness and population dynamics. In both the summer and fall, we observed a negative effect of increasing conspecific larval density on juvenile survival (i.e., the proportion of adult females emerging). However, this trend was much stronger during the fall trial (Figure 1a). This could be due to a hurricane that occurred at the beginning of the fall trial (day three) when larvae were in their most vulnerable stage (i.e., first instar) and/or due to cooler fall temperatures being below the thermal optimum for *Ae. albopictus* larval survival (∼25°C: Mordecai et al., 2017). For example, mean temperatures in the fall were <15°C for three consecutive days. These temperatures, which can result in high *Ae. albopictus* larval and pupal mortality (reviewed in Waldock et al., 2013), combined with the additional stress of developing under high intra-specific competition, could synergistically increase juvenile mortality. This interactive effect of temperature and density on juvenile survival is further supported by our microclimate analyses, which demonstrated that effects of cooler temperatures were more severe at higher densities (Figure 2a).

Contrary to our initial hypothesis and many prior studies (e.g., Ezeakacha and Yee, 2019; Sauers et al., 2022), larvae appeared to develop faster with higher conspecific densities in the cooler fall (Figures 1b, 2b). Several factors could explain this result. First, the timing of juvenile mortality is known to affect the amount of resources available to the surviving larvae (Evans et al., 2022; McIntire and Juliano, 2018; Neale and Juliano, 2019). If mortality of early instar larvae was elevated by cooler fall temperatures and high intra-specific densities, the surviving larvae would have experienced reduced intra-specific competition and increased access to food resources, both of which allow larvae to develop faster (Evans et al., 2022; Neale and Juliano, 2019; Teng and Apperson, 2000). Stage-specific overcompensation has been detected in *Ae. albopictus* when early larval mortality occurred (McIntire and Juliano, 2018). Another possible explanation may be survival bias or frailty. Very few females survived in the fall from the medium-density treatment (7.2%) and even fewer from the high-density treatment (1.2%). If the few individuals that did survive were of higher quality (e.g., competitive ability), then their subsequent development time might be expected to be greater than observed in the low-density treatment. A third explanation could be that larvae facultatively alter their development rate based on a stressor such as low temperatures and/or low quantity of nutritional resources due to increased competition. While not demonstrated before in *Ae. albopictus*, previous work in the *Ae. triseriatus* and *Ae. communis* systems have shown larvae can speed up development rates in environments subject to evaporation (Chodorowski, 1969; Juliano and Stoffregen, 1994).

Our results contrast with the extensive literature documenting relationships between mosquito wing size (i.e. size upon emergence) with variation in conspecific density (Alto et al., 2005; Evans et al., 2021; Lord, 1998) or temperature (Kingsolver and Huey, 2008; Rueda et al., 1990). Warmer temperatures have been shown to typically result in smaller wings and body size – “hotter is smaller”– and cooler temperatures typically result in larger wings and body size – “cooler is bigger” hypothesis (Alto and Juliano, 2001; Ezeakacha and Yee, 2019; Kingsolver and Huey, 2008). Our results contradict these hypotheses. Instead, we consistently found that wing length increased with an increase in the minimum temperature, although the slope of this relationship was less for mosquitoes developing at high conspecific densities (Figure 2c). Only a few studies have demonstrated smaller wing lengths at cooler temperatures (Lounibos et al., 2002; Reiskind and Zarrabi, 2012). This result could be attributed to slower microbial growth in the fall (leading to less food for the developing larvae) or to lower mosquito anabolism (due to cooler temperatures) (Reiskind and Janairo, 2018). Thus, with females in the warmer summer experiencing more microbially rich habitats regardless of larval density treatment as well as a greater capacity to incorporate nutrients into tissue, facilitating more rapid adult female emergence and larger body sizes in the summer. Finally, like development time, mosquito size upon emergence converges at high densities regardless of season, suggesting that high amounts of intra-specific competition could eventually outweigh the influence of variation in abiotic factors such as temperature and relative humidity.

In general, while growth rates declined as larval conspecific densities increased (see also: Edgerly and Livdahl, 1992; Lord, 1998; Reiskind and Lounibos, 2009) and increased at higher temperatures (Alto and Juliano, 2001), we found the estimated per capita growth rates at higher temperatures (e.g., in the summer) were less affected by intra-specific competition than at lower temperatures (e.g., in the fall). This intriguing result suggests that mosquito populations might be less strongly regulated by density-dependent processes as temperatures increase on the landscape. Most studies to date have considered the effects of temperature or conspecific density on growth rates in isolation, without considering the possible interaction. Those that have considered the interaction between temperature and intra-specific competition on per capita growth rates have generally found intra-specific competition to intensify as temperatures warm (Evans et al., 2021; Teng and Apperson, 2000), which is not what we observed in this study. However, we did not account for possible effects of temperature and relative humidity on adult maturation time, longevity, or size-specific fecundity, each of which could potentially modify our estimates of the per capita population growth. Given the interest in vector-pathogen dynamics in response to global change, interactions between temperature and density dependence warrant further investigation.

## CONCLUSION

This study has several important implications. Our study demonstrates that small-scale variation in biotic factors (e.g., intra-specific competition) and abiotic factors (e.g., temperature and relative humidity) are both influential. Importantly, the biotic and abiotic factors interact to affect mosquito population dynamics at local scales. Ignoring the interaction between variation in biotic and abiotic variables could reduce the accuracy and precision of models (no matter their scale) used to predict species distributions and population dynamics, as well as the transmission dynamics of mosquito-borne pathogens. By not accounting for these interactions, mismatches in model predictions could result in unexpected spatial and temporal variation in species densities and host-parasite interactions, especially for organisms with complex life cycles like mosquitoes (Moller-Jacobs et al., 2014; Shapiro et al., 2016). While our results demonstrate the critical role of abiotic factors like temperature and relative humidity that are linked to variation in land use and season, future work should incorporate other aspects of land use and seasonality, such as habitat quality that arises through variation in detrital inputs and precipitation, which will affect the strength of intraspecific competition and potentially the interaction with abiotic variables.

## ACKNOWLEDGMENTS

We thank members of the Murdock lab (UGA and Cornell), Osenberg lab (UGA), and Drake lab (UGA) for their helpful discussions and field support. We particularly thank Sydney Habegger, Luis Hernandez, and Abigail LeCroy for their assistance in the lab and field, Juliana Hoyos, Maria Luisa Müller Theissen, and Kimberly Archbold for their feedback on the Spanish abstract, Ashutosh Pathak for the thoughtful discussions and guidance with the statistical analyses, the property owners who provided us access to the study sites, and several reviewers for helpful comments on previous submissions. We also appreciate the financial support from the National Science Foundation Research Experiences for Undergraduates (Grant No: 1659683), the National Science Foundation Research Traineeship: Interdisciplinary Disease Ecology Across Scales (Grant No: 1545433), the National Institutes of Health Office of Research Infrastructure Programs (Grant No: 2T35OD010433-11), and start-up resources provided by the University of Georgia.

## AUTHORSHIP STATEMENT

CCM designed the experiment. NS, ECH, CWH, PMN, AMS, and JWW conducted fieldwork. NS led, and GRJ assisted with the statistical analyses, with guidance from CCM and CWO. NS drafted the manuscript with guidance from CCM and CWO. All authors were involved in reviewing and editing the manuscript. All authors read and approved the final manuscript.

## DATA AVAILABILITY STATEMENT

The data and code (Solano et al. 2026) that support the findings of this study are openly available in Figshare at https://doi.org/10.6084/m9.figshare.25804639.

