## Appendix for "Mosquito population dynamics are shaped by interactions among larval density, temperature, and relative humidity"

**Journal**:

*Ecology*

**Title**:

**Contents:**

MICROCLIMATE CHARACTERIZATION

*Methods*

*Results*

**Table S1**

**Table S2**

**Figure S1**

**Figure S2**

**Figure S3**

SUPPLEMENTAL FIGURES AND TABLES

**Table S3**

**Table S4**

**Figure S4**

REFERENCES

### MICROCLIMATE CHARACTERIZATION

*Methods.* Following previous work (Evans et al. 2018; Murdock et al. 2017), we used an impervious surface and vegetation cover map to select nine 30 x 30 m sites – three each with low (0-5%), intermediate (6-40%), and high (41-100%) areal cover of impervious surfaces, which we refer to as rural, suburban, and urban land use types respectively (Figure S1).

*Results.* We observed significant main effects of season and land use on microclimate variables (Table S1). Summer temperatures (i.e., mean daily $T_{min}$, $T_{mean}$, and $T_{max}$) were higher than fall temperatures (Figure S2, Table S2). Relative humidity (mean daily $R_{min}$, $R_{mean}$, and $R_{max}$) was also greater in the summer than fall (Figure S2, Table S2). Seasonal variation in diurnal ranges for each abiotic factor also occurred. Mosquitoes in the summer experienced lower fluctuations throughout the day in temperature ($T_{range}$) and relative humidity ($R_{range}$) than in the fall (Table S2). Finally, we observed significant effects of land use on diurnal temperature and relative humidity metrics (Table S1). Urban sites had significantly warmer daily average minimum temperatures ($T_{min}$) than rural sites overall (Table S2) and were less humid than suburban and rural sites (i.e., lower $R_{min}$, $R_{mean}$, and $R_{max}$: Table S2). As a result, urban sites experienced a wider diurnal range in relative humidity ($R_{range}$) than rural sites (Table S2).

In addition to the main effects of season and land use on mosquito microclimate variables, we observed a significant interaction between season and land use on mean daily $T_{max}$ and $T_{range}$ (Table S1). $T_{max}$ was lowest in suburban sites in the summer, but highest in the fall, compared to rural and urban sites (Table S2). Urban sites also had marginally lower average diurnal temperature ranges ($T_{range}$) than rural sites, but only in the fall season (Table S2). We observed an interactive effect of season and land use on three of four metrics of relative humidity: $R_{mean}$, $R_{min}$, $R_{range}$ (Table S1).

Of the eight microclimate variables, several were highly correlated with one another (Figure S3) and one ($R_{max}$) exhibited almost no temporal or spatial variation (Figure S2). We chose three ($R_{min}$, $T_{min}$ and $T_{max}$) that were of a priori interest and were not strongly correlated with one another. These three variables were used to capture variation in the microclimate and potentially explain the observed variation in mosquito traits.

**Table S1.** Type III analysis of variance results from Generalized Linear Mixed Effects Models testing the effects of season (summer vs. fall), land use type (rural, suburban, urban) and their interaction on daily average temperature (ºC) and relative humidity (%). Experimental sites, loggers, and days were included as random factors in all models. χ2=Wald chi-squared statistic; df=degrees of freedom; P=P-value from the Wald test.

| | Daily Minimum | | | Daily Mean | | | Daily Maximum | | | Daily Range | | |
|---|---|---|---|---|---|---|---|---|---|---|---|---|
| | χ2 | df | P | χ2 | df | P | χ2 | df | P-value | χ2 | df | P |
| ***Temperature*** | | | | | | | | | | | | |
| Season | 60.25 | 1 | **<0.001** | 50.91 | 1 | **<0.001** | 17.84 | 1 | **<0.001** | 7.87 | 1 | **0.005** |
| Land use | 8.64 | 2 | **0.013** | 3.98 | 2 | 0.137 | 0.06 | 2 | 0.971 | 1.34 | 2 | 0.512 |
| Season x Land use | 3.16 | 2 | 0.206 | 1.76 | 2 | 0.414 | 7.65 | 2 | **0.022** | 6.19 | 2 | **0.045** |
| ***Relative Humidity*** | | | | | | | | | | | | |
| Season | 1.11 | 1 | 0.292 | 2.09 | 1 | 0.149 | 1.96 | 1 | 0.162 | 2.36 | 1 | 0.125 |
| Land use | 12.37 | 2 | **0.002** | 13.89 | 2 | **0.001** | 9.23 | 2 | **0.01** | 9.99 | 2 | **0.007** |
| Season x Land use | 6.08 | 2 | **0.048** | 8.78 | 2 | **0.012** | 2.43 | 2 | 0.297 | 7.39 | 2 | **0.025** |

**Table S2.** Daily mean microclimate values (95% confidence intervals) across seasons (summer and fall) and land use type (rural, suburban, urban).

| Temperature | $T_{min}$ | | $T_{mean}$ | | $T_{max}$ | | $T_{range}$ | |
|---|---|---|---|---|---|---|---|---|
| (°C) | Summer | Fall | Summer | Fall | Summer | Fall | Summer | Fall |
| | 22.08 | 17.61 | 24.78 | 21.09 | 28.49 | 25.61 | 6.41 | 8 |
| Rural | (21.96, 22.21) | (17.26, 17.96) | (24.63, 24.92) | (20.79, 21.4) | (28.23, 28.76) | (25.28, 25.95) | (6.17, 6.65) | (7.7, 8.3) |
| | 22.34 | 18.27 | 24.78 | 21.39 | 27.91 | 26.58 | 5.57 | 8.31 |
| Suburban | (22.2, 22.47) | (17.88, 18.65) | (24.61, 24.94) | (21.04, 21.73) | (27.66, 28.16) | (26.06, 27.09) | (5.37, 5.78) | (7.83, 8.79) |
| | 22.91 | 18.49 | 25.52 | 21.68 | 29.01 | 25.18 | 6.1 | 6.69 |
| Urban | (22.77, 23.04) | (18.12, 18.85) | (25.36, 25.67) | (21.36, 22) | (28.78, 29.25) | (24.85, 25.5) | (5.92, 6.29) | (6.41, 6.97) |
| Relative | $R_{min}$ | | $R_{mean}$ | | $R_{max}$ | | $R_{range}$ | |
| Humidity (%) | Summer | Fall | Summer | Fall | Summer | Fall | Summer | Fall |
| | 80.5 | 71.31 | 95.87 | 92.28 | 99.9 | 99.56 | 19.4 | 28.25 |
| Rural | (79.41, 81.59) | (69.98, 72.64) | (95.53, 96.22) | (91.77, 92.79) | (99.86, 99.94) | (99.4, 99.72) | (18.32, 20.48) | (26.94, 29.56) |
| | 75.23 | 69.15 | 93.46 | 89.94 | 99.9 | 99.2 | 24.66 | 30.06 |
| Suburban | (74.14, 76.33) | (67.57, 70.72) | (93.03, 93.89) | (89.18, 90.7) | (99.84, 99.96) | (98.92, 99.49) | (23.58, 25.75) | (28.55, 31.57) |
| | 68.07 | 62.24 | 88.53 | 84.7 | 98.82 | 97.29 | 30.74 | 35.04 |
| Urban | (67.05, 69.1) | (60.72, 63.77) | (87.99, 89.07) | (83.82, 85.58) | (98.63, 99) | (96.81, 97.76) | (29.77, 31.72) | (33.64, 36.45) |

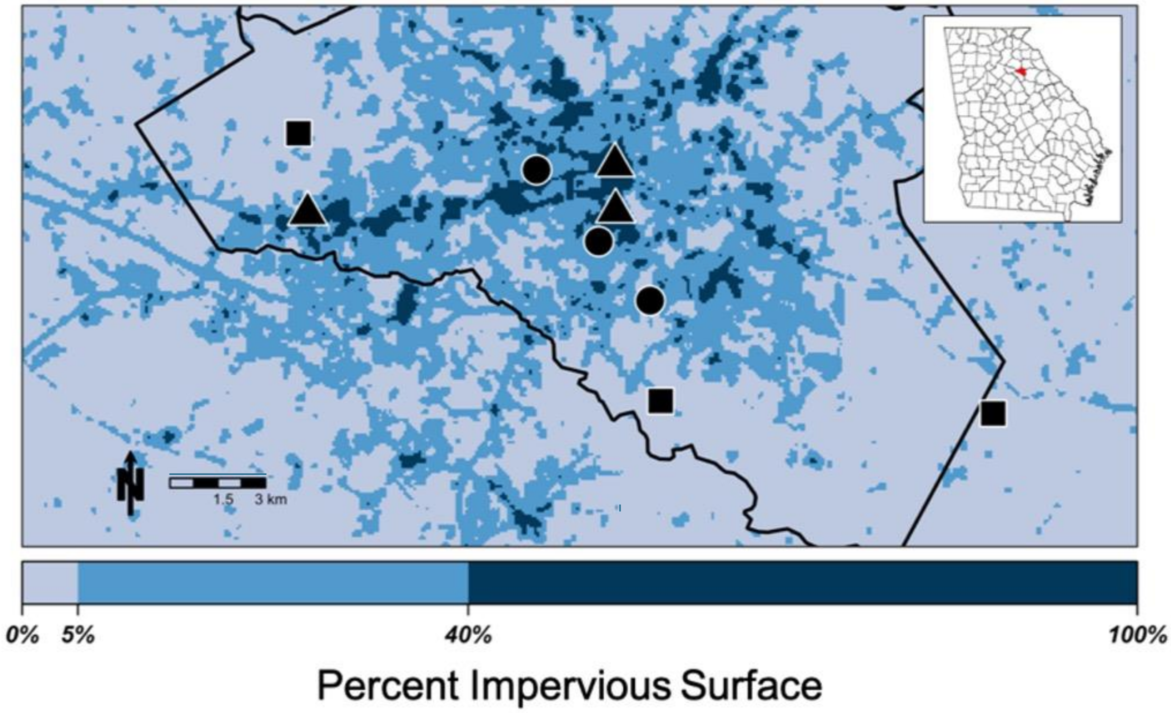


**Figure S1.** Map showing the nine study sites in Athens, GA. Symbols represent land classes (rural=square, suburban=circle, urban=triangle). Color shading represents the amount of impervious surface within 210m of the center of each pixel, as illustrated by the color bar on the bottom, and corresponding to the three land classes (rural: 0-5%; suburban: 5-40%; urban: >40%). Athens-Clarke County is outlined in black and its location within the state is shown in red in the inset map of Georgia.

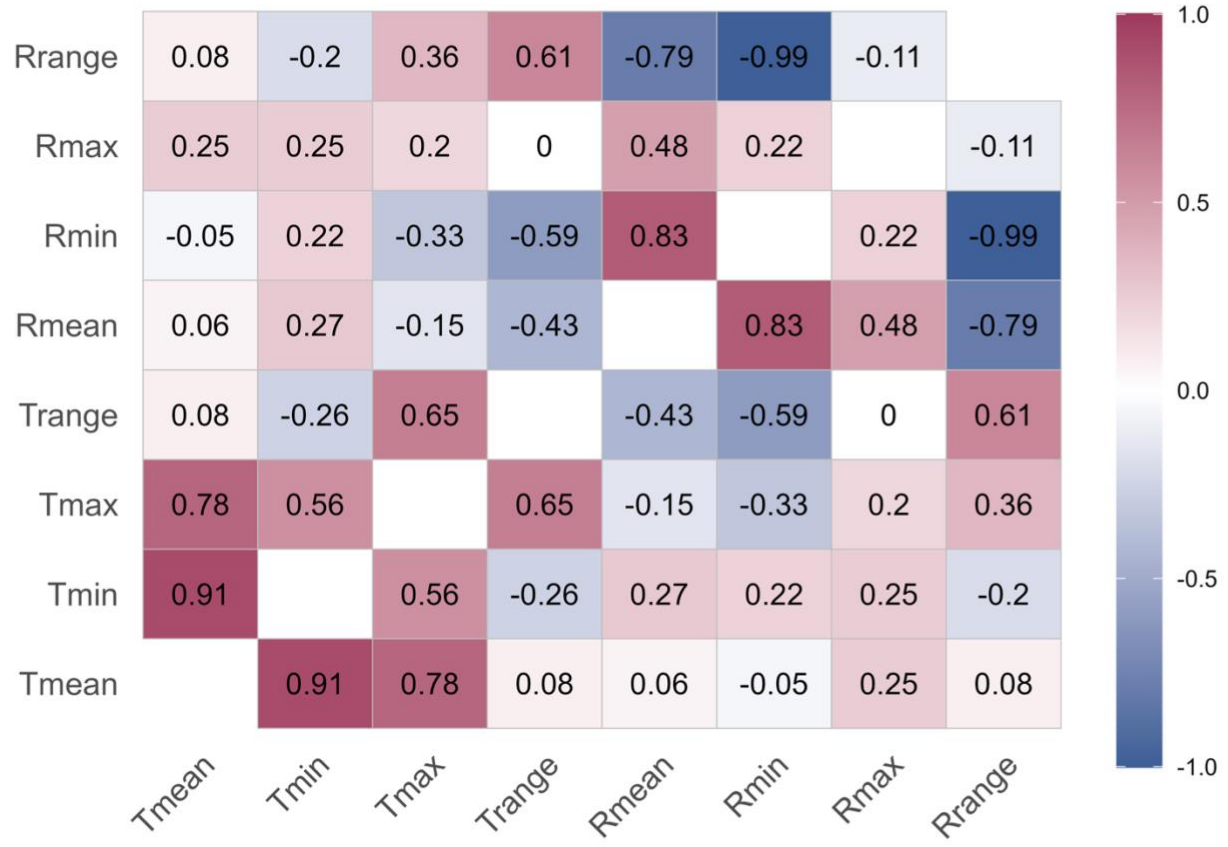


**Figure S2.** Correlogram of microclimate variables giving Pearson's correlation coefficients. Cells are colored according to the magnitude and sign of the correlation coefficient. Note that $R_{max}$ exhibited low correlations with other variables because it was usually 100% and therefore was relatively uninformative (Figure S3d).

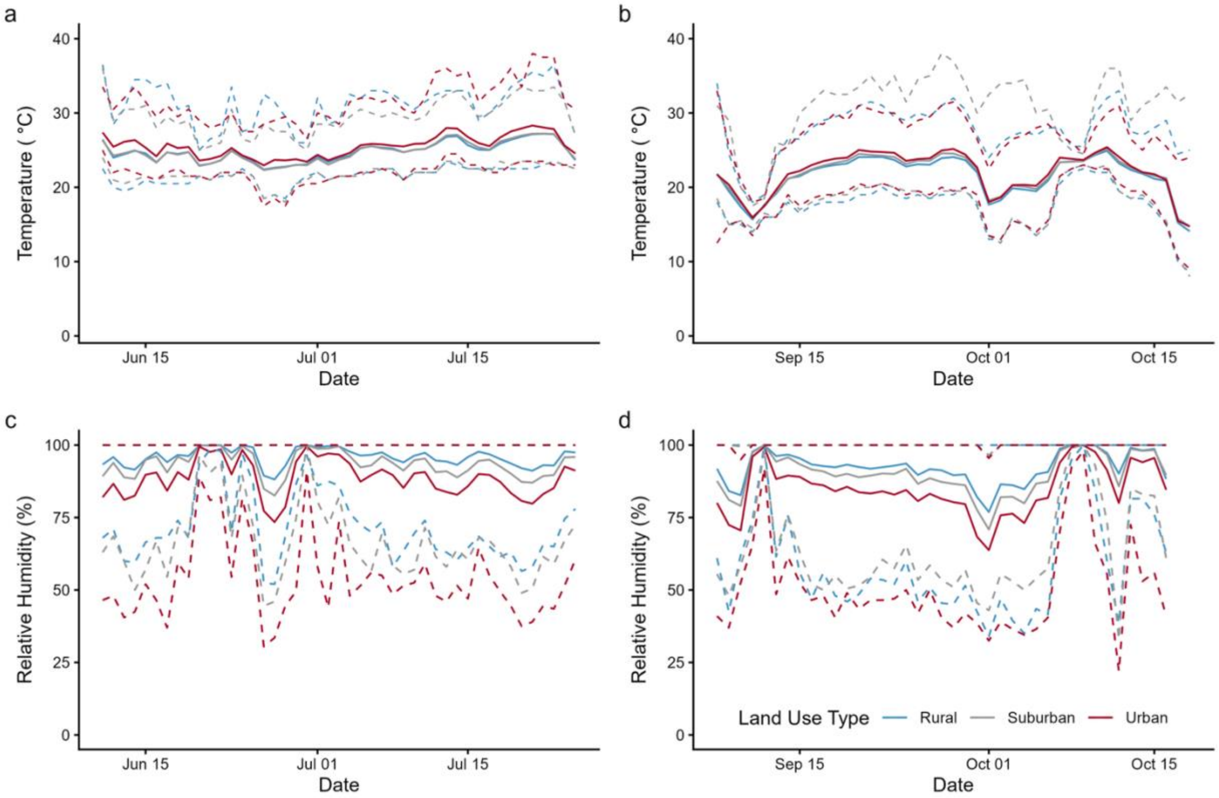


**Figure S3.** Water temperature (ºC) (top panels: a,b) and relative humidity (%) (bottom panels: c,d) at three sites in each of three land use types (rural = blue, suburban = gray, and urban = red) in two seasons [summer 2017 (left panels: a,c) and fall 2017 (right panels: b,d)]. Mean microclimates across the three land use types are in solid lines, with daily maxima and minima indicated by the dashed lines.

### SUPPLEMENTAL FIGURES AND TABLES

**Table S3.** Type III analysis of variance results from Generalized Linear Mixed Effects Models evaluating effects on mosquito traits using the Null, Base, and Microclimate models. The best model(s) are highlighted in blue (see also Table 1 in main text), and significant terms within those models are in **bold**. Factor abbreviations are: S=Season; L=Land use; D=Density; $R_{min}$=mean minimum daily relative humidity, $T_{min}$=mean minimum daily temperature; $T_{max}$=mean maximum daily temperature; $\chi 2$=Wald chi-squared statistic; df=degrees of freedom; P=P-value from the Wald test.

| Model | Emergence (%) | | | | Development Time (days) | | | | Wing Length (mm) | | | | Per Capita Growth Rate, r' ($day^{-1}$) | | | |
|---|---|---|---|---|---|---|---|---|---|---|---|---|---|---|---|---|
| | Factor | $\chi 2$ | df | P | Factor | $\chi 2$ | df | P | Factor | $\chi 2$ | df | P | Factor | $\chi 2$ | df | P |
| Null | D | 407 | 2 | <0.001 | D | 8.9 | 2 | 0.011 | D | 94.4 | 2 | 3E-21 | D | 245 | 2 | ~0 |
| Base | **S** | **228** | **1** | **<0.001** | S | 45.4 | 1 | <0.001 | S | 110 | 1 | <0.001 | **S** | **676** | **1** | **<0.001** |
| | L | 1.2 | 2 | 0.562 | L | 0.24 | 2 | 0.887 | L | 0.07 | 2 | 0.965 | L | 1.66 | 2 | 0.436 |
| | **D** | **425** | **2** | **<0.001** | D | 8.72 | 2 | 0.013 | D | 55.7 | 2 | <0.001 | **D** | **720** | **2** | **<0.001** |
| | SxL | 0.5 | 2 | 0.794 | SxL | 1.59 | 2 | 0.451 | SxL | 5.33 | 2 | 0.07 | SxL | 2.31 | 2 | 0.315 |
| | **SxD** | **186** | **2** | **<0.001** | SxD | 22.9 | 2 | <0.001 | SxD | 6.69 | 2 | 0.035 | **SxD** | **198** | **2** | **<0.001** |
| | LxD | 8 | 4 | 0.09 | LxD | 2.47 | 4 | 0.65 | LxD | 5.25 | 4 | 0.262 | LxD | 4.19 | 4 | 0.381 |
| | SxLxD | 4.1 | 4 | 0.392 | SxLxD | 2.5 | 4 | 0.644 | SxLxD | 6.35 | 4 | 0.174 | SxLxD | 6.13 | 4 | 0.19 |
| Micro | D | 439 | 2 | <0.001 | **D** | **6.26** | **2** | **0.044** | **D** | **88.8** | **2** | **<0.001** | **D** | **631** | **2** | **<0.001** |
| | $T_{min}$ | 56.3 | 1 | <0.001 | **$T_{min}$** | **11.7** | **1** | **<0.001** | **$T_{min}$** | **28.1** | **1** | **<0.001** | **$T_{min}$** | **129** | **1** | **<0.001** |
| | $T_{max}$ | 0.1 | 1 | 0.776 | $T_{max}$ | 3.54 | 1 | 0.06 | $T_{max}$ | 0.87 | 1 | 0.352 | **$T_{max}$** | **25.1** | **1** | **<0.001** |
| | $R_{min}$ | 14 | 1 | <0.001 | $R_{min}$ | 2.92 | 1 | 0.088 | **$R_{min}$** | **30.3** | **1** | **<0.001** | $R_{min}$ | 0.31 | 1 | 0.575 |
| | $DxT_{min}$ | 57.8 | 2 | <0.001 | **$DxT_{min}$** | **8.03** | **2** | **0.018** | $DxT_{min}$ | 0.44 | 2 | 0.801 | **$DxT_{min}$** | **46.1** | **2** | **<0.001** |
| | $DxT_{max}$ | 8.6 | 2 | 0.014 | $DxT_{max}$ | 2.11 | 2 | 0.349 | $DxT_{max}$ | 4.56 | 2 | 0.102 | **$DxT_{max}$** | **23.9** | **2** | **<0.001** |
| | $DxR_{min}$ | 74.1 | 2 | <0.001 | $DxR_{min}$ | 1.69 | 2 | 0.43 | **$DxR_{min}$** | **6.24** | **2** | **0.044** | $DxR_{min}$ | 0.98 | 2 | 0.612 |

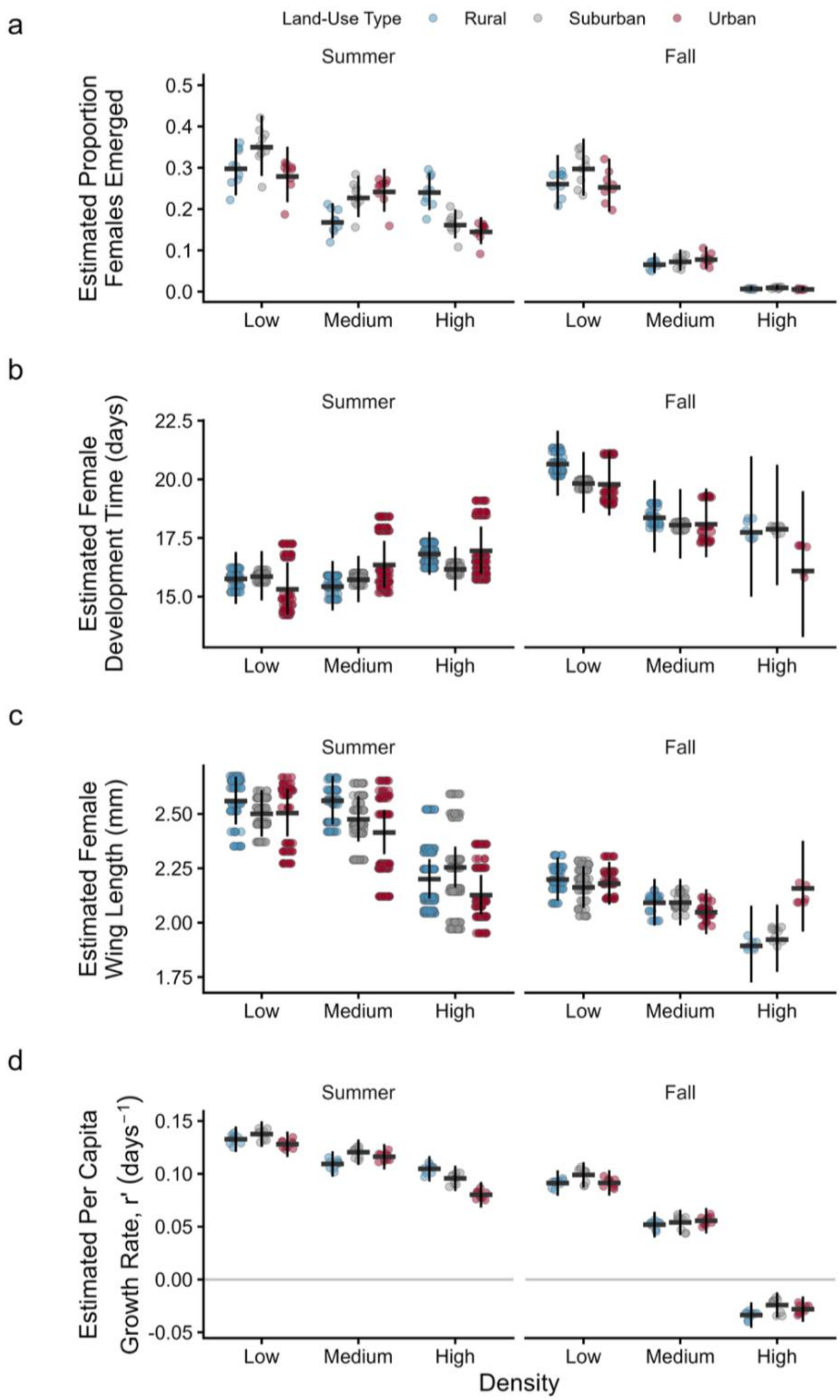


**Figure S4.** Mixed effects model predictions (mean +/- 95% CI) of the demographic mosquito traits of *Ae. albopictus:* (**a**) the proportion of female larvae that emerged (i.e., survival probability), (**b**) mean female *Ae. albopictus* development time (days), (**c**) mean female wing length (mm), and (**d**) per capita growth rate, $r'$ (day$^{-1}$), for each larval density treatment (30 = low, 60 = medium, 120 = high) across each land-use type (rural = blue, suburban = gray, urban = red). Points are for individual females (b,c) or jars (a,d) adjusted for random effects.
